# Soma-germ contact across basement membrane in ovary

**DOI:** 10.1101/2024.07.10.602885

**Authors:** Yasuhiko Chikami, Kensuke Yahata

## Abstract

Epithelial cells generally interact with other cells and environments at their apical side. At the basal surface, the basement membrane impedes such an interaction. One notable instance is communication between soma and germ cells within the ovary across numerous bilaterian taxa. This contact underlies proper oogenesis and subsequent embryogenesis through facilitating nutrient supply to gametes, exchange of molecules and ions, and participation in signal transduction for primary axis formation. Throughout layers of the history of morphology and cellular biology, there has been emphasis on this heterocellular interaction primarily occurring at the apical side of epithelial cells. Contrary to this long-standing understanding, we uncover that ovarian follicle cells in three myriapod species belonging to phylogenetically basal myriapod clades extend their cytoplasmic processes, penetrating the basement membrane to establish direct contact with oocytes. These discoveries demonstrate the ovarian soma-germ interaction transverse the basement membrane, suggesting that the basal matrix does not always a physical barrier to soma-germ communication. Furthermore, we find that the ovarian somatic cells in a myriapod directly connect with the oogonia or young oocyte before the formation of their basement membrane. This result suggests an overlooked construction manner of heterocellular communication at the basal domain of epithelial cells.

## Introduction

Epithelium, a cellular layer covering animal organs, commonly exhibits polarity along the apicobasal axis (Buckley and Johnston, 2022; Eaton and Simons, 1995; Gibson and Perrimon, 2003) Along the basal domain, these cells are consistently bordered by a layer of basement membrane—an extracellular matrix serving as a barrier between the epithelium and adjacent tissues (Glentis et al., 2014; Jayadev and Sherwood, 2017; LeBleu et al., 2007; Stanley et al., 1982). In contrast, the apical domain of most cells is free from such a matrix and features microvilli or cytoplasmic projections to face the external environment directly, including coelomic fluid and neighboring tissues (Apodaca, 2018; Kloc and Kubiak, 2017; Matsumoto et al., 1987). Consequently, contacts and communications between the epithelial and other cells often occur at the apical surface.

One prominent example of epithelial cell contact with other cells is soma-germ communication in the ovary. Female somatic cells near germ cells, referred to as accessory, follicle, or granulosa cells, frequently establish physical connections with developing oocytes in various bilaterians (Buccione et al., 1990; Büning, 1994; Eckelbarger, 2005; Eckelbarger and Hodgson, 2021; Hadgson and Eckelbarger, 2000; Rouse and Tzetlin, 1997). This contact underlies proper oogenesis and subsequent embryogenesis through facilitating nutrient supply to gametes, exchange of molecules and ions, and participation in signal transduction for primary axis formation (Anderson and Woodruff, 2001; Bohrmann and Haas-Assenbaum, 1993; Brooks and Woodruff, 2004; Clarke, 2018; Giorgi and Postlethwait, 1985; Huebner, 1981; Kloc and Kubiak, 2017; Munz, 1988; Telfer and Woodruff, 2002; Waksmonski and Woodruff, 2002). The polarized epithelial cells face the oocytes at their apical side. Consistent with the general canon, these polarized cells contact germ cells at their apical side through their cytoplasmic projections and heterocellular junctions.

In the context of the positional relationship between the epithelial polarity and oocytes, the ovarian structure of Myriapoda holds significance. The ovarian structure in Myriapoda is characterized by the basement membrane between the follicle epithelium and the oocyte, with oocyte growth occurring within a hemocoelic space—an extraovarian environment (figure 1a) (Chikami and Yahata, 2019, 2024; Kubrakiewicz, 1991a; Miyachi and Yahata, 2012; Yahata et al., 2018). Therefore, the follicle cells face oocytes at their basal side. There have been no reports of ovarian soma-germ connections in Myriapoda despite extensive information on ovarian morphology in numerous species over the past century (Biliński, 1979; Chikami and Yahata, 2019, 2024; Fontanetti et al., 2010; Herbaut, 1974; Jangi, 1957; Knoll, 1974; Kubrakiewicz, 1991a, 1991b; Miyachi and Yahata, 2012; Sareen and Adiyodi, 1983; Tiegs, 1940, 1945, 1947a, 1947b; Tönniges, 1902; Yahata et al., 2018; Yahata and Makioka, 1994, 1997; Yahata, 2012). Furthermore, it was observed that a diplopod species lacks the heterocellular connection (Kubrakiewicz, 1991a). These studies suggest that the basement membrane serves as a barrier against the soma-germ interaction in accordance with the general cannon.

**Figure 1.**
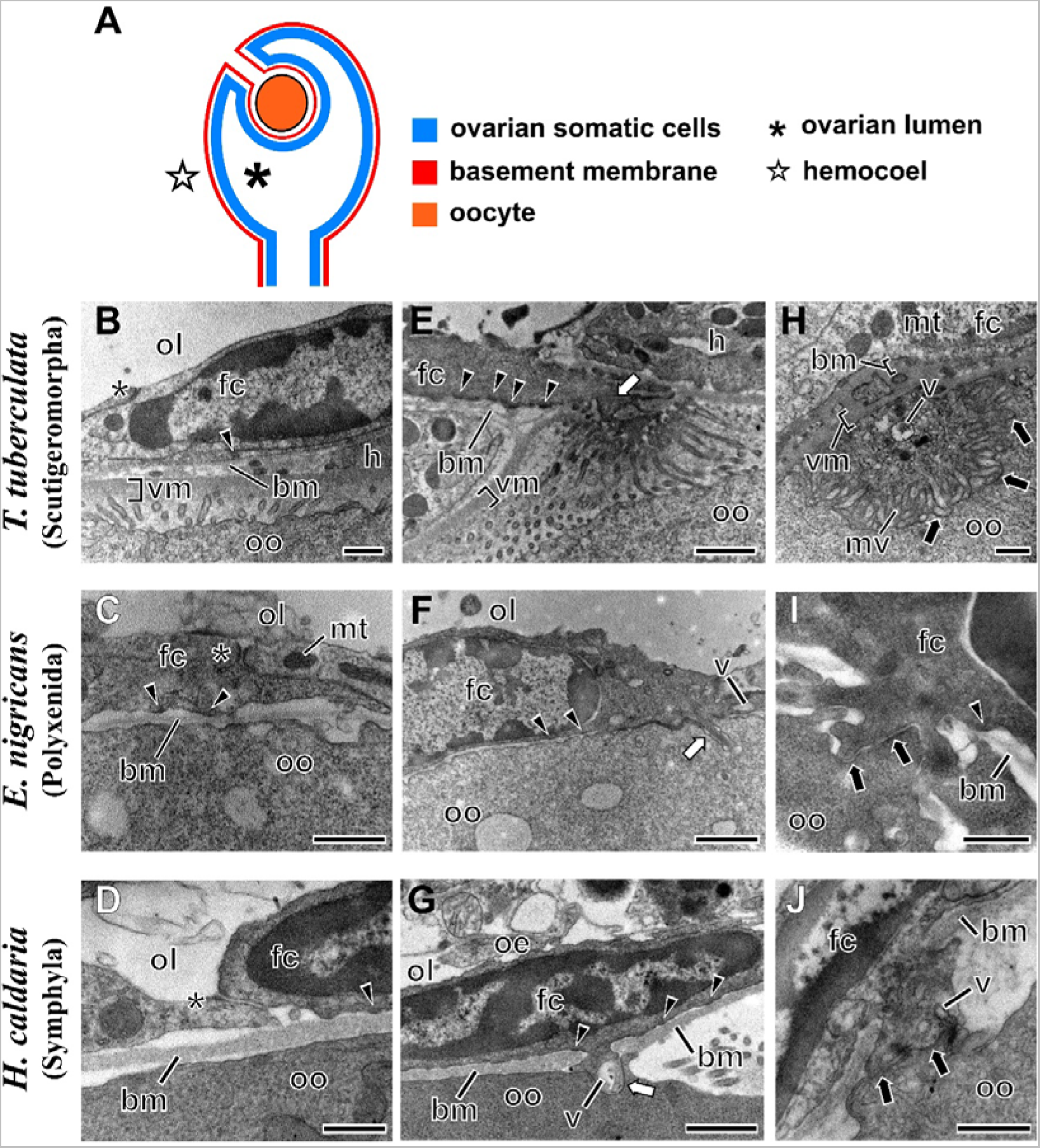
Ovarian structure of Myriapoda and ultrastructural nature of the follicle cells. (A) Schematic image of the Myriapoda-type ovary based on Miyachi and Yahata (2012). The basic morphology of the follicle epithelium in *T. tuberculata* (B), *E. nigricans* (C), and *H. caldaria* (D). The follicle cells extending cytoplasmic processes (white arrows) in *T. tuberculata* (E), *E. nigricans* (F), and *H. caldaria* (G). The soma-germ contact in *T. tuberculata* (H), *E. nigricans* (I), and *H. caldaria* (J). Note that the cytoplasmic process in (H) was largely interwoven, resulting in an image that is not continuous with the follicle cells. Asterisks, arrowheads and black arrows indicate the adherens junction, hemidesmosomes and heterocellular junction, respectively. bm: basement membrane, fc: follicle cell, h: hemocyte, mt: mitochondria, mv: microvilli, oe: ovarian epithelium, ol: ovarian lumen, oo: oocyte, v: vesicle. Scales: 1 µm (E–G), 0.5 µm (B–D, H–J).

In contrast to such a conventional knowledge, here, we unexpectedly uncover that follicle cells contact oocytes across the basement membrane in *Thereuonema tuberculata* and *Eudigraphis nigricans*, representatives of the most basal centipede and millipede clades, respectively (Benavides et al., 2023; Blanke and Wesener, 2014; Edgecombe and Giribet, 2004; Fernández et al., 2016, 2018; Miyazawa et al., 2014; Rehm et al., 2014; Rodriguez et al., 2018), and *Hanseniella caldaria*, a member of Symphyla, the most basal myriapod clade in certain phylogenetic inferences (Miyazawa et al., 2014; Rehm et al., 2014). In a broader sense, this finding suggests novel perspectives on ovarian soma-germ communication and unique modes of heterocellular interaction.

## Materials and Methods

### Animals

*Hanseniella caldaria* (Hansen, 1903) (Symphyla: Scutigerellidae) adults were collected in Tsuchiura City, Ibaraki Prefecture, and Ibusuki City, Kagoshima City, Japan, between May 2015 and March 2016. *Thereuonema tuberculata* (Wood, 1862) (Chilopoda: Scutigeromorpha: Scutigeridae) adults were collected in Sakuragawa City, Ibaraki Prefecture, Japan, between January and October 2016. *Eudigraphis nigricans* (Miyosi, 1947) (Diplopoda: Polyxenida: Polyxenidae) adults were collected in Hitachinaka City, Ibaraki Prefecture and Sagamihara City, Kanagawa Prefecture, Japan, between June and October 2016. Some individuals were kept in plastic cases at 25°C.

### Transmission electron microscopy

In *H. caldaria*, the whole body including the ovary was cut into some blocks of 1 mm in length. In *T. tuberculata*, and *E. nigricans*, the ovary was dissected from the body and sectioned. Each sample was pre-fixed with 2.5% glutaraldehyde in 0.1 M phosphate buffer (PB) (pH 7.0–7.2) for 3 hours at room temperature and then post-fixed with 1% Osmium tetroxide (OsO_4_) in the same buffer for 1 hour at 4°C. After washing with PB, the fixed samples were dehydrated in a graded acetone series and then immersed and embedded in the Quetol-651 resin (Nisshin EM, Tokyo, Japan). The resin blocks containing the samples were polymerized for 36–60 hours at 60°C and then were cut into ultrathin sections of 50–80 nm with a diamond knife on an ultramicrotome (EM UC7, Leica Microsystems, Wetzlar, Germany). The sections where then viewed under a transmission electron microscopy (H-7650, Hitachi, Tokyo, Japan) equipped with a charge-coupled device camera (Velta 2k×2k, Olympus, Tokyo, Japan) at 80 kV.

## Results

First, to confirm the Myriapoda-type ovarian structure in investigated species, we observed the follicle cells at ultrastructural level. The follicle epithelium in all investigated species consisted of a single layer of flattened cells (Figure 1B–D). The follicle cells were lined with a homogenous mono-laminated matrix, consistent with features of a basement membrane described in previous studies of ovaries in myriapods, including *H. caldaria*^27–30^. This basement membrane was situated between the cells and oocytes (Figure 1B–D). The thickness of the basement membrane was massive in *H. caldaria* (Figure 1D) but was quite thin in *E. nigricans* (Figure 1C) and *T. tuberculata* (Figure 1B). Electron-dense structures, like hemidesmosomes, were found between the follicle cells and the basement membrane (Figure 1B–G). On the opposite side, adjacent cells adhered to each other through adherens junctions (asterisks in Figure 1B–D). Thus, the follicle cells faced oocytes at their basal domain.

We then elaborately observed the space between oocytes and follicle cells to explore a soma-germ connection. The follicle cells of *T. tuberculata*, *E. nigricans*, and *H. caldaria* extended their cytoplasmic processes to the oocyte (Figure 1E–G). These processes exhibited varying shapes among the species, with a smooth shape in *E. nigricans* (Figure 1F) and *H. caldaria* (Figure 1G) and a microvilli-bearing structure in *T. tuberculata* (Figure 1E). Notably, they penetrated the basement membrane of the follicle epithelium, reaching the surface of the oocyte in all species. The cytoplasmic processes bordered on the oocyte with narrow gaps (Figure 1H–J). Electron-dense materials were also distributed along the gaps. These features of the contact zone consisted with those of a gap junction. Therefore, in all three myriapods, the cytoplasmic process of the follicle epithelium was directly in contact with the oocyte through heterocellular junctions. Furthermore, some vesicular structures existed in the follicle cells contacting with oocytes (Figure 1F, G, H, J).

Finally, to elucidate how ovarian somatic cells pierce the basal matrix in Myriapoda, we observed the contact zone between the follicle cells and oocytes during oogenesis in *T. tuberculata*. We found that somatic cells closely contacted germ cells, which were either oogonia or young oocytes (Figure 2A, B). At this stage, we could not detect a basement membrane in the somatic cells, indicating that the cells were unpolarized. During the previtellogenic stage, the polarized follicle cells extended cytoplasmic processes across the basement membrane (Figure 2C). These cytoplasmic processes persisted during the vitellogenic stage, penetrating the basement membrane and thick double-layered vitelline membranes (Figure 2D). Therefore, the soma-germ contact occurred before the basement membrane formation was completed and existed throughout the oogenesis in *T. tuberculata*.

**Figure 2.**
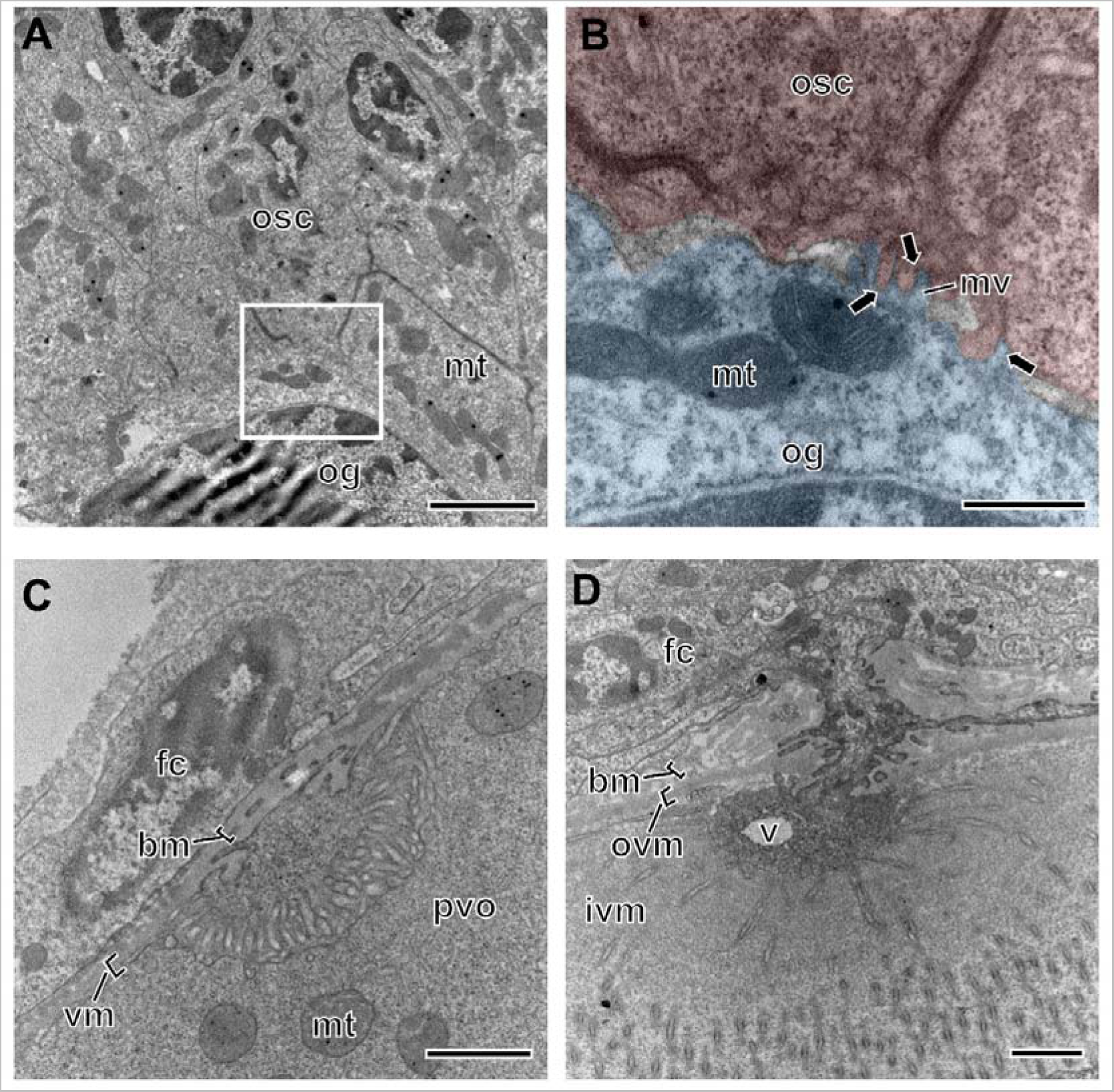
The closely contact between germ cells and somatic cells during oogenesis. (A) The oogonia stage. (B) The enlargement view of the white frame of (A). The blue and red areas indicate the oogonia and the ovarian somatic cell, respectively. (C) The previtellogenic stage. (D) The vitellogenic stage. Black arrows indicate the soma-germ connection. bm: basement membrane, fc: follicle cell, ivm: inner vitelline membrane, mt: mitochondria, mv: microvilli, og: oogonia, oo: oocyte, osc: ovarian somatic cell, ovm: outer vitelline membrane, v: vesicle. Scales: 2 µm (A), 0.5 µm (B), 1 μm (C, D).

## Discussion

We demonstrated that the cytoplasmic processes of follicle cells penetrate the basement membrane, establishing connections with oocytes in three myriapods: *H. caldaria*, *T. tuberculata*, and *E. nigricans*. Close contacts between female germ cells and the cytoplasmic processes or microvilli of ovarian somatic cells have been observed across various ecdysozoan groups, including Nematoda (Hall et al., 1999; Velde and Coomans, 1988), Chelicerata (Alberti and Palacios-Vargas, 2015; Biliński et al., 2008; Dumont and Anderson, 1967; Jędrzejowska et al., 2014; Miyazaki and Biliński, 2006; Oliveira et al., 2005; Ōsaki, 1971; Talarico et al., 2009; Witaliński and Žuwała, 1981), and Pancrustacea (Biliński, 1987; Biliński and Klag, 1982; Biliński et al., 1985; Biliński and Szklarzewiez, 1987; Huebner, 1981; Jaglarz et al., 2014; Mazzini and Giorge, 1985; Souty, 1980; Tsutsumi et al., 2005). However, Myriapoda has traditionally been considered an exception because such contact has not been previously described, in addition to a previous report indicating the absence of heterocellular connections in a diplopod species (Kubrakiewicz, 1991a). Furthermore, certain features of the myriapod ovary, such as a persistent basement membrane between follicle cells and germ cells,^26–28^ have supported this interpretation. Contrary to this conventional belief, our findings represent evidence of soma-germ communication in Myriapoda, highlighting the ubiquity of soma-germ contact among Arthropoda.

More broadly, the contact between follicle cells and oocytes in myriapods challenges a traditional canon in soma-germ connections within the ovary. In the ovaries of many animals, including vertebrates, nematodes, and insects, epithelial cells typically adhere to developing oocytes (Buccione et al., 1990; Büning, 1994; Eckelbarger and Hodgson, 2021; Glentis et al., 2014; Hadgson and Eckelbarger, 2000; Hall et al., 1999; Rouse and Tzetlin, 1997). These cells face oocytes at their apical side, establishing soma-germ interactions through heterocellular junctions. To our knowledge, there have been no reports of ovarian somatic cells contacting oocytes by penetrating their basement membrane. Our findings provide the first evidence of ovarian soma-germ interactions across the basement membrane, suggesting that the basement membrane does not always serve as a barrier between epithelial cells and other tissues, including germ cells.

In certain tissues, epithelial cells can communicate with other cells at the basal side through some distinct manners (Apodaca, 2018; Rowe and Weiss, 2008). During some developmental processes, e.g., epithelial-mesenchymal transition and oncogenesis, epithelial cells sometimes undergo de-epithelization and break their basement membrane (Kelley et al., 2014; Morrissey et al., 2013). Also, cells near the epithelium, such as neural, muscle, and immune cells, can insert projections like tunneling nanotubules into epithelial cells through pores in the basement membrane (Apodaca, 2018; Naphade et al., 2015). These two ways are common in that de-epithelialized or non-epithelial cells breach the basement membrane. Additionally, it is hypothesized that epithelial cells and adjacent cells burrow filopodia-like thin projections, cytonemes, into the basement membrane, establishing communication with each other in the matrix (Gonzalez-Mendez et al., 2017, 2019; Gradilla et al., 2018). Therefore, epithelial cells can interact with other cells at their basal surface through loss of epitheliality, invasion by other-type cells, or utilizing the basement membrane as a scaffold rather than crossing it. By contrast, the follicle cells in the myriapod ovary keep their epithelial nature, extend their cytoplasmic process, and fully penetrate the basement membrane. In this context, it is suggested that the soma-germ interaction in the myriapod ovary likely represents a simple but overlooked mode of heterocellular interaction across the basement membrane.

The soma-germ contact in *T. tuberculata* was established before the formation of the basal matrix. While further examination is needed to determine whether the earliest contacts are homologous to those after the previtellogenic stage, this result suggests that the penetration of the basement membrane may be a secondary consequence of basal matrix secretion around the soma-germ contact rather than a result of the breakdown of the completed matrix. If so, the construction of the soma-germ contact in *T. tuberculata* differs from the previously known heterocellular connections at the basal domain that occur after basement membrane secretion (Apodaca, 2018; Kelley et al., 2014; Morrissey et al., 2013; Naphade et al., 2015). In this study, we could not observe the polarization process of the follicle cells or the construction of the soma-germ connection in the other two species due to the developmental stages of the samples. Further investigation into the mechanism and manner of soma-germ contact across the basement membrane in myriapod species is needed to test our concept.

The species in focus of this study belong to the basal centipede and millipede groups^46–53^ and Symphyla. Moreover, the soma-germ connection is generally involved in somatic cell contribution to oogenesis (Glentis et al., 2014; Matova and Cooley, 2001). Consequently, it is inferred that the ovary with the soma-germ connection persisted in the common ancestor of Myriapoda, engaging in communication with oocytes and potentially playing a role in oogenesis. This inference supports that the absence of the ovarian soma-germ contact reported in myriapods may be a secondary loss during the myriapod evolution, which could be a unique evolutionary polarity in Myriapoda. Further observations, including taxa yet to be examined, will enhance this evolutionary history.

Overall, our findings unveil a more widespread prevalence of ovarian soma-germ contact among arthropods than previously believed and present evidence for ovarian soma-germ interaction across the basement membrane, an exception to the conventional view in zoology. The myriapod ovary may be a goldmine of discoveries that can be exceptions to conventional canons.

## Acknowledgements

We thank Mr. K. Kojima, Mr. R. Nagasawa, Mr. N. Naya, Mr. G. Takahashi, Ms. Y. Takatani, Ms. M. Shibata, and Ms. E. Umetani for their encouragement to collect myriapods. Our analysis of transmission electron microscope was carried out with instruments at the Open Facility, Research Facility Center for Science and Technology, University of Tsukuba.

## Author contributions

Yasuhiko Chikami, Conceptualization, Methodology, Investigation, Resources, Writing – Original Draft, Visualization. Kensuke Yahata, Conceptualization, Methodology, Resources, Writing – Review & Editing

